# Blood-brain Barrier Crossing using Magnetic Stimulated Nanoparticles

**DOI:** 10.1101/2021.12.23.472846

**Authors:** Jingfan Chen, Muzhaozi Yuan, Caitlin A Madison, Shoshana Eitan, Ya Wang

**Affiliations:** J. Mike Walker ‘66 Department of Mechanical Engineering, Texas A&M University, College Station, TX 77843; Department of Psychological and Brain Sciences, Texas A&M University, College Station, TX, 77843; Department of Electrical and Computer Engineering, Texas A&M University, College Station, TX, 77843; Department of Biomedical Engineering, Texas A&M University, College Station, TX, 77843

**Keywords:** PBPK modeling, Magnetic nanoparticles, BBB crossing

## Abstract

Due to the low permeability and high selectivity of the blood-brain barrier (BBB), existing brain therapeutic technologies are limited by the inefficient BBB crossing of conventional drugs. Magnetic nanoparticles (MNPs) have shown great potential as nano-carriers for efficient BBB crossing under the external static magnetic field (SMF). To quantify the impact of SMF on MNPs’ *in vivo* dynamics towards BBB crossing, we developed a physiologically based pharmacokinetic (PBPK) model for intraperitoneal (IP) injected superparamagnetic iron oxide nanoparticles coated by gold and conjugated with poly(ethylene glycol) (PEG) (SPIO-Au-PEG NPs) in mice. Unlike most reported PBPK models that ignore brain permeability, we first obtained the brain permeabilities with and without SMF by determining the concentration of SPIO-Au-PEG NPs in the cerebral blood and brain tissue. This concentration in the brain was simulated by the advection-diffusion equations and was numerically solved in COMSOL Multiphysics. The results from the PBPK model after incorporating the brain permeability showed a good agreement (regression coefficient R^2^ = 0.825) with the *in vivo* results, verifying the capability of using the proposed PBPK model to predict the *in vivo* biodistribution of SPIO-Au-PEG NPs under the exposure to SMF. Furthermore, the *in vivo* results revealed that the brain bioavailability under the exposure to SMF (4.01%) is slightly better than the control group (3.68%). In addition, the modification of SPIO-Au-PEG NPs with insulin (SPIO-Au-PEG-insulin) showed an improvement of the brain bioavailability by 24.47 % in comparison to the non-insulin group. With the SMF stimulation, the brain bioavailability of SPIO-Au-PEG-insulin was further improved by 3.91 % compared to the group without SMF.

## 1. Introduction

Brain therapeutics face many challenges, one of which is the low permeability and high selectivity of the blood brain barrier (BBB) to conventional drugs [1]. The BBB, which covers the surface area of cerebrovascular capillaries, is composed of a single layer of endothelial cells that are joined by tight junctions and end-feet of astrocytes. This structure prevents solutes in the circulating blood from non-selectively crossing into the extracellular fluid of the CNS [2, 3]. Nanoparticle (NP)-based drug-carrier systems have been reported to aid in retention and specific delivery of a multitude of potential therapeutic agents across the BBB due to their excellent tunable surface functionality [4-8]. Among them, magnetic nanoparticles (MNPs) earned interest because of their unique properties to respond to magnetic fields, which greatly improve the targeting efficiency [9-15]. Superparamagnetic iron oxide (SPIO) NPs are small synthetic γ-Fe_2_O_3_, Fe_3_O_4_ or α-Fe_2_O_3_ particles with a core ranging from 10 nm to 100 nm in diameter, which have a unique property of superparamagnetism that can be guided to a specific tissue or organ by an external magnetic field [16]. However, uncoated SPIO NPs have several disadvantages, such as tending to aggregate [17] and causing cellular toxicity [18]. One way to overcome this is to coat them using biocompatible materials, such as gold (Au), which exhibits excellent biocompatibility, low toxicity, and surface functionalization because of its inertness and stability [19]. In our previous work, we have demonstrated that SPIO coated by Au (SPIO-Au) NPs exhibit excellent biocompatibility, excellent cellular uptake ability, and a promotional effect of neuronal growth and differentiation on PC-12 and primary neuron cells [20]. However, NPs as foreign objects are readily removed from systemic circulation by macrophages, impeding accumulation in the targeted tissue. To promote retention and BBB crossing efficiency, coating the surface of NPs with poly (ethylene glycol) (PEG) can prevent the NPs from aggregation, opsonization, and phagocytosis, thereby prolonging circulation time [21-24]. In addition, modifying NPs with ligands, such as insulin [25] and transferrin [26], can facilitate the BBB crossing by the receptor-mediated endocytosis process.

To get insights into the efficiency of magnetic targeting and the brain accumulation of MNPs, *in vivo* studies still stand as a fundamental complementary step. Researchers have demonstrated the effective and direct passage of MNPs across the BBB and accumulating in the brain site near the magnet in animals under an external static magnetic field (SMF) [6, 11]. However, characterization of the transport process for MNPs through a mathematical model is crucial for elucidating their transport across BBB, especially for understanding how blood circulating MNPs interact with the cerebral microvasculature under exposure to an external SMF. Current models studying MNPs’ targeted delivery are limited to leaky tumor vasculature (without considering the complexity of the BBB) or local vessel network (without considering the whole-body circulation), which cannot precisely reflect the behavior of NPs in brain [27-30]. Thus, it is crucial to have a comprehensive quantitative model that can predict the kinetics of MNPs in different tissues/organs and estimate the amount that reaches the brain. Recently, researchers have employed physiologically based pharmacokinetic (PBPK) models to describe the *in vivo* biodistribution of NPs [31-34]. PBPK models were first used in pharmaceutical research and drug development to estimate the internal dose of toxic agents or their metabolites in target tissues [35]. In PBPK models, the pharmacokinetic behavior of a chemical substance in the body-that is, its absorption, distribution, metabolism, and elimination (ADME)-is represented by equations that attempt to quantitatively describe actual physiological processes [35, 36]. However, so far, the currently developed PBPK models cannot precisely predict the accumulation of NP in the brain as they usually assume zero brain permeability.

In this paper, we developed a PBPK model with the ADME process shown in Fig.1 to estimate the *in vivo* biodistribution of intraperitoneal (IP) injected PEG coated SPIO-Au NPs (SPIO-Au-PEG) in mice under an exposure to SMF. We first estimated the brain permeabilities in the absence of magnetic field and under the exposure to SMF by simulating the behaviors of SPIO-Au-PEG NPs in the brain using the advection-diffusion equations. Other pharmacokinetic parameters of the NPs in the PBPK model were then calibrated with the *in vivo* data from the control group. The developed PBPK model adequately predicted the biodistribution of SPIO-Au-PEG NPs under the exposure of SMF, indicating the feasibility of simulating the *in vivo* biodistribution of MNPs. In addition, we compared the accumulation of SPIO-Au-PEG NPs in the brain under no magnetic field (MF-), the SMF, and the dynamic magnetic field (DMF). We also explored the strategy based on insulin modified SPIO-Au-PEG (SPIO-Au-PEG-insulin) NPs crossing the BBB and demonstrated that insulin modification and SMF stimulation can enhance the BBB crossing of SPIO-Au-PEG NPs.

## 2. Material and Methods

### 2.1 Synthesis of SPIO-Au NPs

The SPIO-Au NPs were synthesized according to our previously reported method [20]. In detail, 2 mL of 4.66 mM SPIO (Ferrotec EMG 304) solution was stirred with 6 mL of 0.1 M sodium citrate for 10 min to enable the exchange of the absorbed OH^-^ with citrate anions. The mixture was then diluted to 100 mL with deionized (DI) water. 0.5 mL of 1% HAuCl_4_ solution was added to the mixture. The pH was adjusted to 9-10 by using 0.1 M NaOH solution. Then 0.6 mL of 0.2 M NH_2_OH·HCl was added to the mixture to form the Au coating. The color of the mixture changed from brown to purple in several minutes. After that, another 0.5 mL of 1% HAuCl_4_ was added to the solution, followed by the addition of 0.2 mL of 0.2 M NH_2_OH·HCl. This process was repeated several times to form a thicker Au coating. The color of the final solution changed from purple to red. Chemicals used in this section were purchased from Sigma-Aldrich (St. Louis, MO).

### 2.2 Functionalization of SPIO-Au NPs with PEG (SPIO-Au-PEG)

PEG5000 is commonly used to greatly enhance the blood circulation time of NPs [37]. In this paper, we prepared two sets of SPIO-Au-PEG NPs to use throughout this study. For the first set, 45 uL of 5 mg/mL SH-mPEG (MW∼5 kDa) was added to 0.785 mL of SPIO-Au NPs at the concentration of 1.274 mg/mL. This first set of SPIO-Au-PEG1 NPs was used to investigate the effect of different magnetic fields. For the second set, 45 uL of 5 mg/mL PEG solution was composed by a mixture of 85% SH-mPEG (MW∼5 kDa) and 15% SH-PEG-COOH (MW∼3.4 kDa) and was added to 0.785 mL of SPIO-Au NPs at the concentration of 1.274 mg/mL. This second set of SPIO-Au-PEG2 NPs was used to exam the effect of insulin modification. The resulting PEG-coated SPIO-Au NPs were collected by centrifugation at 12,000 rpm for 15 minutes and washed twice to remove the uncoated PEG. The final concentration of SPIO-Au-PEG NPs was maintained at 3 mg/mL for the *in vivo* test purpose. PEG used in this section were purchased from Biochempeg (Watertown, MA, USA).

### 2.3 Functionalization of SPIO-Au-PEG NPs with insulin (SPIO-Au-PEG-insulin)

To facilitate the BBB crossing by the receptor-mediated endocytosis process [25], the strategy of insulin modification on SPIO-Au-PEG NPs was explored. In detail, 2 mM of 1-Ethyl-3-(3-dimethylaminopropyl) carbodiimide HCl (EDC, 22980-Sigma-Aldrich, St. Louis, MO, USA) and 5 mM of N-hydroxysulfosuccinimide sodium salt (NHS, 24500-Sigma-Aldrich, St. Louis, MO, USA) were added directly to 1 mL of insulin (11061-Sigma-Aldrich, St. Louis, MO, USA) at the concentration of 10 mg/mL. The solution was mixed well and reacted for 15 minutes at room temperature. 0.02 mL of the resulted mixture was then added directly to 12 mL of 1 mg/mL SPIO-Au-PEG2 NPs. The solution was left to stir overnight to ensure the conjugation of the PEG layer to the insulin, and SPIO-Au-PEG-insulin NPs were purified after the solution was centrifuged. The final concentration was 3 mg/mL.

### 2.4 Characterization of SPIO-Au, SPIO-Au-PEG and SPIO-Au-PEG-insulin NPs

Transmission electron microscope (TEM) was used to assess the morphological characteristics and to determine the average core size of SPIO and SPIO-Au NPs. After suspension in DI water and sonication for 10 min to avoid the formation of large aggregates, SPIO and SPIO-Au NPs were placed on a carbon-coated grid, dried, and observed under TEM at 80 kV. The average size of the NPs was calculated from at least 100 particles. The hydrodynamic diameters of SPIO-Au, SPIO-Au-PEG1, SPIO-Au-PEG2 and SPIO-Au-PEG-insulin NPs and their zeta-potential were measured by dynamic light scattering (DLS) with a Zetasizer apparatus (Malvern Instruments; Malvern, UK). The hydrodynamic diameter was calculated from the intensity-weighted distribution function obtained by CONTIN analysis of the correlation function embedded in Malvern software. All measurements were performed in triplicates.

### 2.5 The prediction of biodistribution of SPIO-Au-PEG NPs using PBPK model

With a goal of predicting SPIO-Au-PEG NPs concentration in the brain and other organs, it is necessary to elucidate the route of NPs in the body. In Fig. 1, the ADME processes of SPIO-Au-PEG NPs after IP injection in mice is represented schematically. Firstly, the SPIO-Au-PEG NPs are rapidly absorbed from the peritoneal cavity. After that, there are two possible pathways for SPIO-Au-PEG NPs to reach systemic circulation: (1) they are drained into portal circulation and then into the systemic circulation, or (2) they bypass the liver to get into the systemic circulation directly [38]. At this stage, NPs can be distributed to different organs. Finally, they will be eliminated via metabolism or excretion from the body, where the liver (bile excretion and feces elimination) and the kidney (renal excretion) are the major organs responsible for NPs excretion [39].

**Fig. 1.**
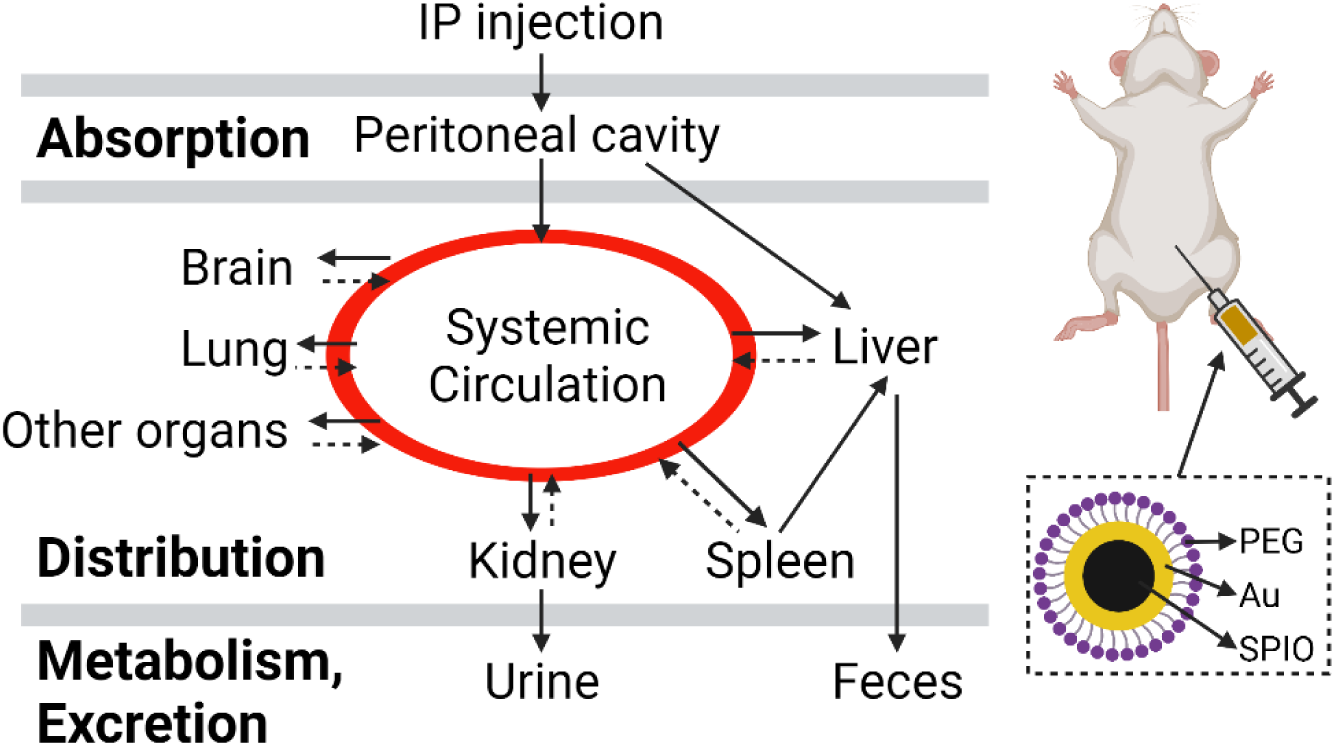
The ADME process of IP injection of SPIO-Au-PEG NPs in mice. The black lines represent confirmed routes for NPs, and the dashed lines represent hypothetical routes. The IP injected SPIO-Au-PEG NPs are rapidly absorbed from the peritoneal cavity, and then distributed to different organs from systemic circulation. Finally, they are eliminated via metabolism or excretion from the body. Created with BioRender.com.

#### 2.5.1 Mathematical description of the PBPK model

The ADME processes are described and analyzed using a whole-body PBPK model consisting of seven compartments: blood, liver, spleen, kidneys, lungs, brain, and remainder as shown in Fig. It is established based on the previously developed PBPK models [40] to quantitatively describe the biodistribution of SPIO-Au-PEG NPs. The PBPK model has the advantage of providing the concentration-time profiles of NPs in individual organs. It can also be used for cross-species extrapolation, which allows scaling-up of the animal data to humans. In this study, based on the macroscopic observation that a portion of the dose remained trapped in the intraperitoneal space, an optimized absorbed amount (76%) of SPIO-Au-PEG NPs getting into the blood circulation is assumed according to the reference that studied the biodistribution of IP injected gold NPs (AuNPs) [41]. Following absorption into the bloodstream, SPIO-Au-PEG NPs were distributed from the systemic circulation to the organs. Subsequently, they passed through the capillary membrane of the organs to be taken up via endocytosis, which was described by the membrane-limited model. The excretion rates of NPs by the liver and kidneys are assumed constant. Therefore, the SPIO-Au-PEG NPs concentrations of all compartments and sub-compartments contained within the PBPK model were mathematically described by a set of ordinary differential equations (ODEs).

The differential equations used to describe the transportation between arterial blood and capillary blood of each organ (systematic circulation) as shown in Fig. 2a:

**Fig. 2.**
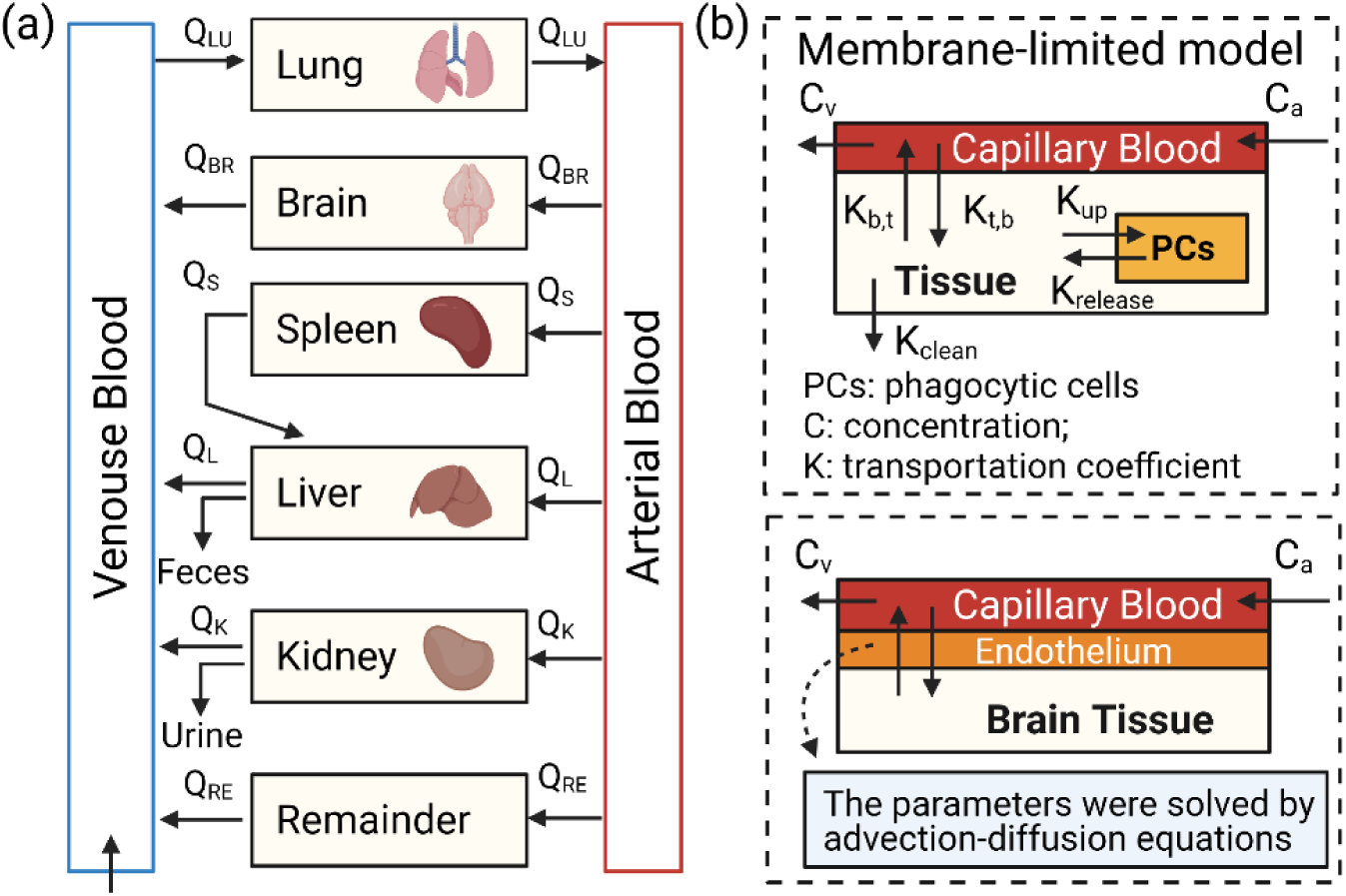
(a) The whole-body PBPK model for SPIO-Au-PEG NPs in mice. Q and C are blood flow and SPIO-Au-PEG NPs concentration for lungs (Lu), brain (BR), spleen (S), liver (L), kidney (K), and remainder (RE). (b) Sub-compartment representation of organs in the PBPK model. PCs were included in organs except for the brain and the remainder. For tissues without PC, K_up_ and K_release_ will be zero; while without elimination, K_clean_ will be zero. C_a_ is the inflowing arterial concentration, C_v_ is the outflowing venous concentration. The brain permeability was obtained from advection-diffusion equations.

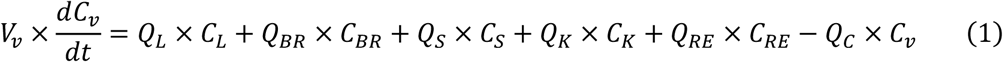

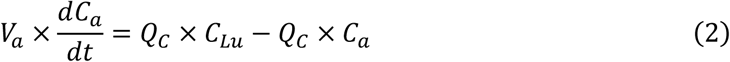

Inside each organ, as shown in Fig. 2b, which contains the capillary space and the tissue space, SPIO-Au-PEG NPs transfer between these two spaces can be expressed as:

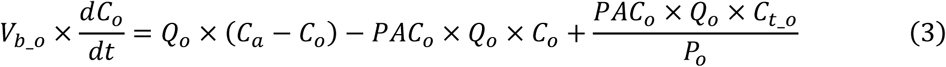

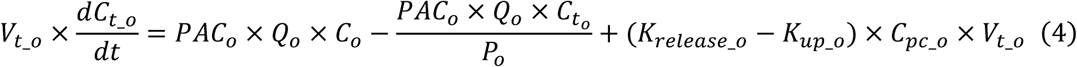

where *C*_*o*_(mg/L), *C*_*t*_*o*_(mg/L) and *C*_*pc*_*o*_are the concentration of SPIO-Au-PEG NPs in the capillary blood of the organ, the tissue of the organ and the phagocytic cells (PCs) of the organ, respectively. *C*_*v*_(mg/L) and *C*_*a*_(mg/L) are the concentration of SPIO-Au-PEG NPs in the venous and arterial blood, respectively. The physiological parameters, *Q*_*o*_(L/h) is the blood flow to the organ; *Q*_*C*_ (L/h) is the cardiac output; *V*_*v*_(L), *V*_*a*_(L), *V*_*b*_*o*_(L) and *V*_*t*_*o*_(L) are the volume of the venous blood, arterial blood, the capillary blood of the organ and the tissue of the organ respectively, and can be obtained from the literature based on the weight of the mice [42]. *PAC*_*o*_(unitless) is the permeability coefficient between capillary blood and tissue, and *P*_*o*_(unitless) is the ratio of tissue and blood distribution coefficients for the organ; *K*_*release*_*o*_(per h) is the release rate constant of SPIO-Au-PEG NPs from PCs to the tissue in the organ, and *K*_*up*_*o*_(per h) is the uptake rate of SPIO-Au-PEG NPs from the tissue to PCs in the organ (it varies with time). SPIO-Au-PEG NPs are assumed to be taken by PCs via endocytosis, which described by the uptake rate as [40]:

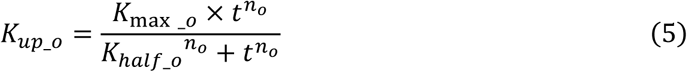

where *t* (h) is the simulation time, *K*_*max*_*o*_(per h) is the maximum uptake rate parameter in the organ, *K*_*half*_*o*_(h) is the time reaching half of *K*_*max*_*o*_, and *n*_*o*_(unitless) is the Hill coefficient. The *o* in the parameter subscript represents lung (Lu), brain (BR), spleen (S), liver (L), kidney (K), and remainder (RE). By solving the above equations, the concentrations of SPIO-Au-PEG NPs in major organs can be obtained.

The mean percentage of blood flow to the organs and the percentage of organ volumes (capillary blood and tissue) to the body weight were adopted from the literature [42]. Given the mice weight of 0.025 kg (the average weight of the mice from our experiment), the physiological parameters including blood flow to different organ (*Q*_*o*_), the organ volume (*V*_*b*_*o*_and *V*_*t*_*o*_) and the blood volume (*V*_*v*_and *V*_*a*_), were calculated and given in Table 1.

**Table 1.** Physiological parameters used in PBPK model.

| Parameter | Symbol | Mean Value |
| --- | --- | --- |
| Body weight (kg) | BW | 0.025 |
| Cardiac output (L/h/kg<sup>0.75</sup>) | QCC | 16.5 |
| **Blood flow to organ (L/h)** |  |  |
| Liver | $Q_L$ | 0.167 |
| Spleen | $Q_S$ | 0.011 |
| Kidneys | $Q_K$ | 0.094 |
| Lungs | $Q_{Lu}$ | 1.037 |
| Brain | $Q_{BR}$ | 0.034 |
| Remainder | $Q_{RE}$ | 0.730 |
| **Volume of the tissue in organ (kg)** |  |  |
| Liver | $V_{t_L}$ | $9.49 \times 10^{-4}$ |
| Spleen | $V_{t_S}$ | $1.04 \times 10^{-5}$ |
| Kidneys | $V_{t_K}$ | $3.23 \times 10^{-4}$ |
| Lungs | $V_{t_{Lu}}$ | $8.75 \times 10^{-5}$ |
| Brain | $V_{t_{BR}}$ | $4.12 \times 10^{-4}$ |
| Remainder | $V_{t_{RE}}$ | $2.09 \times 10^{-2}$ |
| **Volume of the blood (kg)** |  |  |
| Venous Blood | $V_V$ | $9.80 \times 10^{-4}$ |
| Arterial Blood | $V_a$ | $2.45 \times 10^{-4}$ |
| **Volume of the capillary blood in organs (kg)** |  |  |
| Liver | $V_{b\_L}$ | $4.26 \times 10^{-4}$ |
| Spleen | $V_{b\_S}$ | $2.13 \times 10^{-5}$ |
| Kidneys | $V_{b\_K}$ | $1.02 \times 10^{-4}$ |
| Lungs | $V_{b\_Lu}$ | $8.75 \times 10^{-5}$ |
| Brain | $V_{b\_BR}$ | $1.28 \times 10^{-5}$ |
| Remainder | $V_{b\_RE}$ | $8.70 \times 10^{-4}$ |

For the brain compartment, the permeability is solved by simulating the behavior of SPIO-Au-PEG NPs in the brain model that considers the BBB, which will be discussed in next section. While for other compartments, the following pharmacokinetic parameters of SPIO-Au-PEG NPs are fitted based on the *in vivo* data: the distribution coefficients (*P*_*o*_), the permeability coefficients (*PAC*_*o*_), the endocytosis-related parameters (*K*_*max*_*o*_, *K*_*half*_*o*_, *n*_*o*_and *K*_*release*_*o*_), and the elimination rate constants for the kidney (*K*_*urine*_) and liver (*K*_*bile*_). The model structure could then be built with the defined parameters in Matlab R2020 to simulate the biodistribution of SPIO-Au-PEG NPs in mice.

#### 2.5.2 Estimation of brain permeability using advection-diffusion equations

The permeability for the brain compartment is usually assumed to be zero in most reported PBPK models by assuming a highly efficient BBB [43], which cannot correctly reflect the accumulation of NPs in the brain. To estimate the permeability coefficient in the brain, we considered the scenario where SPIO-Au-PEG NPs enter the brain via the circle of Wills (CoW). The CoW is an important junction of arteries that supplies blood to the brain. In our *in vivo* experiments, we employed C57BL/6 mice (8-week, male, 25 g) to collect the experimental results of SPIO-Au-PEG biodistribution. Therefore we reconstructed the CoW based on the mouse brain model from the literature [44], which analyzed the brain vascular features of the same mouse type (12-week, male, 28.9 g. Because the body weight of male mice between 8 weeks and 12 weeks is similar, it is assumed the brain vasculature size is also similar). As shown in Fig. 3a, two pairs of arteries supply blood to the brain: the left and right internal carotid arteries (ICA) and the left and right vertebral arteries (VA). The ICAs divide into the anterior cerebral arteries (ACA) and continue to form the middle cerebral arteries (MCA). And the VAs fuse into the basilar artery (BA), which then branches into the posterior cerebral arteries (PCA). According to the statistical analysis, the total inflow of blood to the brain is distributed to each ICA (36% each) and VA (14% each) [45]. From table 1, the total blood flow to the brain is 0.034 L/h. Thus, the blood flow rates for the inlets (ICA and VA) were calculated to be 0.012 and 0.004 L/h, respectively, which were within the range as reported in the literature [46]. Then the whole blood flow velocity field of the CoW was determined by the known vascular geometry of the mouse and the inlets’ velocity, using the Fluid Flow Module in COMSOL Multiphysics.

**Fig. 3.**
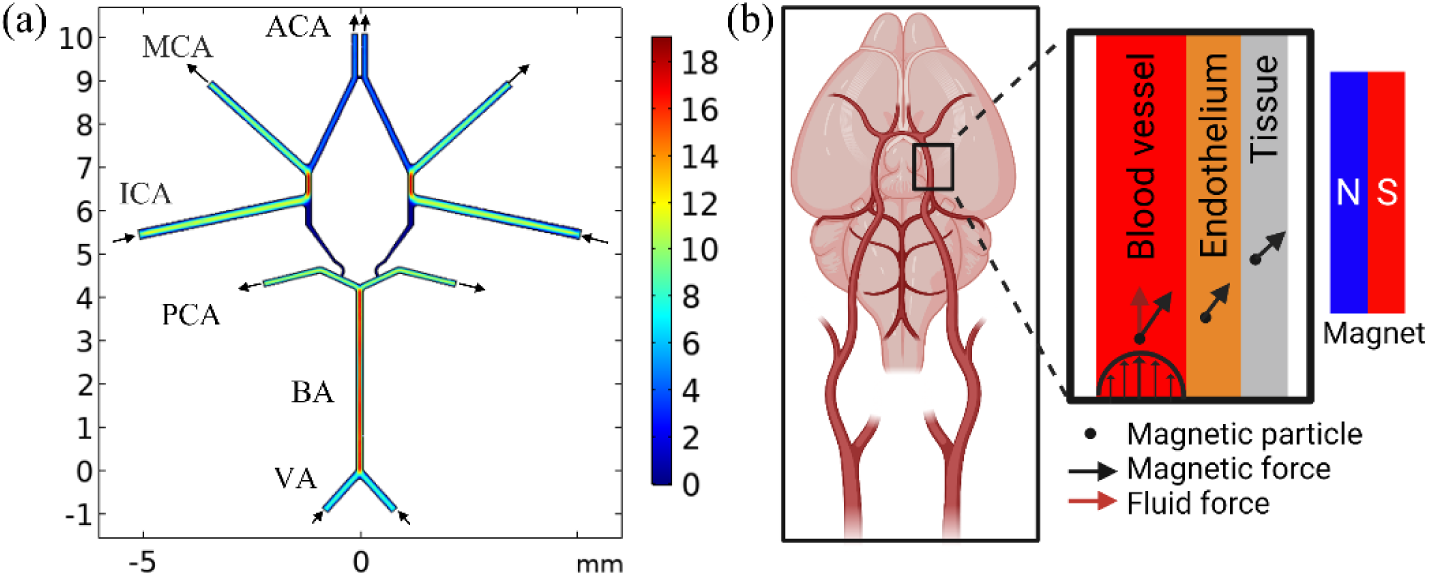
(a) The blood velocity of the CoW of the mouse (color bar: cm/s); (b) the simulated model of SPIO-Au-PEG NPs transport in the cerebral blood and the brain tissue. The NPs within the blood vessel experience diffusion, advection by blood flow, and magnetic forces. The NPs in the surrounding endothelial and tissue layer experience diffusion and magnetic drift but no blood flow forces.

We considered that there are three layers for the CoW model as shown in Fig. 3b: a blood vessel, an endothelial layer and a tissue layer. For particles with a diameter below 40 nm one solves an advection-diffusion equation for a certain particle density, rather than the Newtonian equation for the trajectory of a single particle [47]. Thus, the motion of SPIO-Au-PEG NPs in each layer of the CoW is simulated as an advection–diffusion process for the particle concentration, which is a function of time and is governed by three effects: diffusion, advection by blood flow, and magnetic drift.

Diffusion is driven by a gradient in concentration. Due to the small size of SPIO-Au-PEG NPs, Brownian motion is significant, which causes random movement of particles. The Brownian diffusion D_B_ is given by:

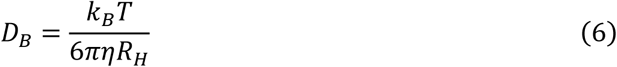

Here *T* is the absolute temperature (measured in Kelvin) and *k*_*B*_ is Boltzmann’s constant. *η* is the dynamic blood viscosity and *R*_*H*_ is the particle hydrodynamic radius. Another diffusive mechanism that influences the NPs motion in vessels is shear-induced diffusion. Blood is a highly concentrated fluid with red blood cells (RBCs) suspended in plasma where sheared cell-cell collisions give rise to random motions with a diffusive character. The shear-induced diffusion coefficient, D_S_ is given by:

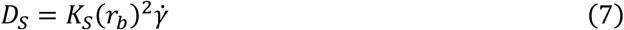

where *K*_*S*_ ≈ 5 × 10^−2^ is a dimensionless coefficient dependent on RBCs concentration, *r*_*b*_ is the radius of RBCs and 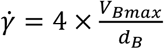 is the local value of the fluid shear rate. *V*_*Bmax*_ and *d*_*b*_ are the maximum centerline velocity and the diameter, respectively. Thus, the total diffusion coefficient in the blood is:

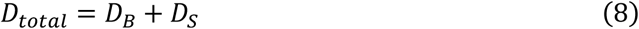

The diffusion coefficient in the endothelial layer is calculated by an empirical equation [48]:

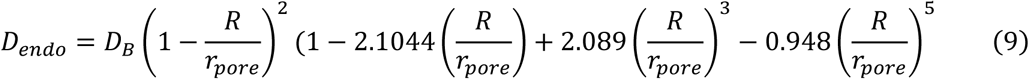

where *R* is the radius of SPIO-Au-PEG NP and *r*_*pore*_ ≈ 10 *nm* is the average radius of the pores in a membrane. While the diffusion coefficient in the tissue is defined as [48]:

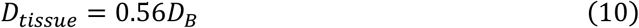

As the blood flows, SPIO-Au-PEG NPs will move along with the velocity of the blood, which triggers the advection of NPs. The advection effect is determined by the blood velocity. Since the blood velocity is highest at the centerline and is almost zero at the walls due to the no-slip boundary condition, SPIO-Au-PEG NPs near the blood vessel wall experience a much smaller advection effect and can potentially be driven by a much smaller magnetic force. The velocity of blood in the CoW was solved in COMSOL as shown in Fig. 3a.

The magnetic force acted on each SPIO-Au-PEG NP is given by [49],

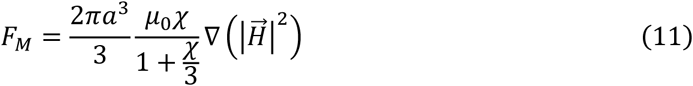

where *a*is the radius of the magnetic core of SPIO-Au-PEG NPs, *χ* is the magnetic susceptibility, *μ*_0_ is the permeability of free space and 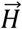 is the magnetic field strength. When the magnetic force is applied to a particle, it will accelerate the particle in the direction of this force until it reaches an equilibrium velocity. The opposing Stokes drag force on a spherical particle is given by,

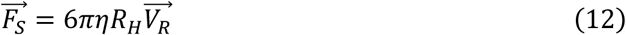

where *R*_*H*_ is the hydrodynamic radius of SPIO-Au-PEG NP and *V*_*R*_ is the relative velocity. When the Stokes drag force first equals the applied magnetic force, the particle will reach its equilibrium relative velocity, defined as,

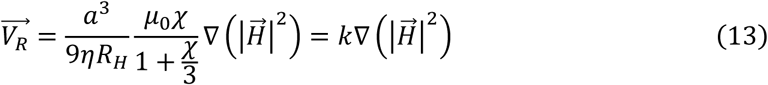

where 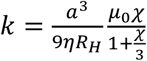 is the magnetic drift coefficient.

Thus, considering these effects together, the concentration of SPIO-Au-PEG NPs in the cerebral blood is given by:

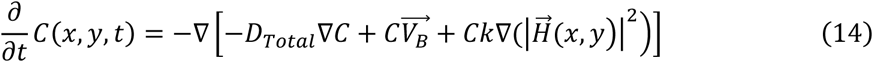

where *C* is the concentration of SPIO-Au-PEG NPs,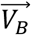 is the blood velocity.

SPIO-Au-PEG NPs in the surrounding endothelial and tissue layer only experience diffusion and magnetic drift but no blood flow forces. The concentration inside the membrane and tissue is defined more simply by the equation:

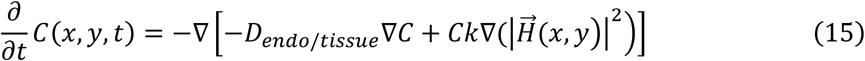

After solving the equations, the permeability coefficient of the brain was calculated as [50]:

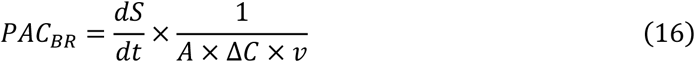

where *dS/dt* is the amount of SPIO-Au-PEG NPs passing the membrane in unit time, *A* is the surface area of the capillaries, Δ*C* is the average concentration difference between the cerebral blood and blood tissue and *v* is the average blood velocity. The permeability coefficients of SPIO-Au-PEG NPs for the brain without and with SMF were further incorporated into the previously described PBPK model to estimate the amount of NPs staying in the brain tissue for every systemic circulation.

### 2.6 In vivo biodistribution of SPIO-Au-PEG and SPIO-Au-PEG-insulin NPs in mice

For the *in vivo* study, mice were IP injected with SPIO-Au-PEG NPs or SPIO-Au-PEG-insulin NPs. IP injection represents a safe method of introducing materials into animals. Given the fast absorption of most substances from the peritoneal cavity, it is generally considered that systemic exposure of the IP-administered substance is an acceptable and justifiable rodent model for intravenous (IV) injection [38]. 8-week-old male C57BL/6 N mice were purchased from Envigo Lab (Houston, Texas, USA) and housed in a temperature-controlled (21±2°C, humidity 45%) vivarium with a 12-hour light/12-hour dark cycle (lights on at 07:30). 92 mice were divided for three tests. (1) For test 1, to study the effect of magnetic fields, mice were IP injected with SPIO-Au-PEG1 NPs at 30 mg/kg and were randomly divided into three treatment groups as follows: (a) MF-, (b) SMF and (c) DMF. The mice were anesthetized and sacrificed at 4, 8 and 12 hr post-injection (n = 4/time-point). (2) For test 2, to study the effect of insulin in the absence of magnetic field, mice were randomly divided into two treatment groups: injected with (a) SPIO-Au-PEG2 NPs (insulin-) and (b) with SPIO-Au-PEG-insulin NPs (insulin+). All NPs were IP injected at 30 mg/kg. The mice were anesthetized and sacrificed at 4, 8 and 12 hr post-injection (n = 4/time-point). (3) For test 3, to study the effect of SMF with the existence of insulin, mice were IP injected with SPIO-Au-PEG-insulin NPs at 30 mg/kg and were randomly divided into two treatment groups as follows: (a) MF- and (b) SMF. The mice were anesthetized and sacrificed at 2, 4, 8 and 12 hr post-injection (n = 4/time-point). At the designated post-injection time, the brain (dissected into midbrain, cortex and cerebellum), liver, lung and blood samples were collected and frozen immediately at −80°C. All procedures were conducted in accordance with the National Institutes of Health Guide for the Care and Use of Laboratory Animals and were approved by the Institutional Animal Care and Use Committee.

#### 2.6.1 Static and dynamic magnetic applicator

To investigate the effect of an externally applied magnetic field on the accumulation of SPIO-Au-PEG NPs in the brain, we firstly constructed a SMF using a circular Halbach array composed of 8 magnets (NdFeB, grade N52, core strength of 1.48 T, dimensions 12.7 × 12.7 × 12.7 mm, K&J Magnetics, Inc.). The magnetization direction of the magnets arranged around the circle (Fig. 4a) provided a strong magnetic field inside and near zero magnetic field outside. The array was specialized for a mouse’s head as a ring with 16 mm thickness, 38 mm inner diameter (approximately the size of the diameter of the mouse head) and 78 mm outer diameter. Specifically, to generate a DMF, a DC motor controlled by L298N motor driver, and an Arduino board was used to generate a rotation speed at 60 rpm on this array (Fig. 4a). This applicator was used for test 1.

**Fig. 4.**
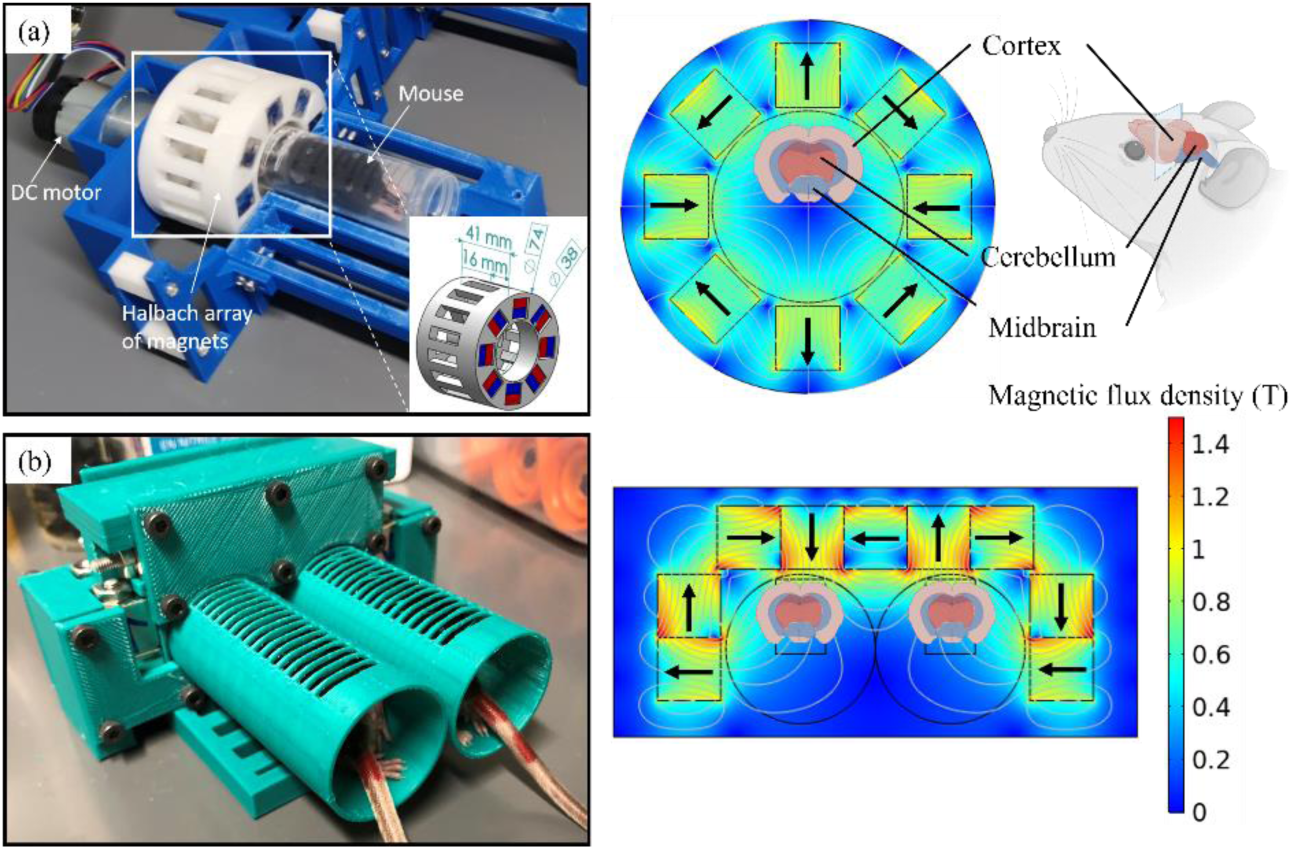
Left, *In-vivo* animal experimental set up and right, the magnetic flux diagram (simulated from COMSOL) of (a) the circular Halbach array of magnets for SMF and DMF (rotation at 60 rpm) and (b) the linear Halbach array. Different brain sections located inside the applicator are labelled (cortex has the strongest magnetic exposure).

To increase the strength of SMF around the head of the mice, a linear Halbach array composed of 9 magnets for treating two mice simultaneously was designed (Fig. 4b). The magnetization direction of the magnets is shown in Fig. 4b. This applicator was used for test 3. For all mice, the magnetic applicator was applied around their head immediately after IP injection of NPs for 1 hour. Mice’s heads were re-exposed to the magnetic applicator every fourth hour for 1 hour until euthanized.

#### 2.6.2 Au distribution determination by ICP-MS spectrometry

The Au level can reflect the distribution of SPIO-Au-PEG or SPIO-Au-PEG-insulin NPs. In order to quantitatively investigate the amount of SPIO-Au-PEG or SPIO-Au-PEG-insulin NPs in major organs, the Inductively Coupled Plasma Mass Spectrometer (ICP-MS) technique was used to determine the content of Au. Tissue samples were dried at 60°C for 24 h r and then all samples were pre-digested with 5 mL ∼65% nitric acid (TraceMetal^™^ Grade, Thermo Fisher Scientific, Waltham, MA) and 1 mL ∼30% hydrochloric acid (TraceMetal^™^ Grade, Thermo Fisher Scientific, Waltham, MA) overnight. Samples were then heated in Digiprep MS digestion block (SCP Science, Champlain, NY, USA), followed by the addition of 2 mL ∼30% hydrogen peroxide (16911-Sigma-Aldrich, St. Louis, MO, USA). After the digestion, the samples were diluted into 5% nitric acid and 1% hydrochloric acid and tested on the PerkinElmer NexION 300D (PerkinElmer, Inc.). A calibration curve with known Au concentrations was prepared and the Au concentration was determined according to absorbance values, compared to calibration curves. The results are shown as mean values.

All results are represented as the average value ± standard error of the mean (SEM) based on at least 3 biological replicates. Due to the small sample size (n=4/time point), statistical analysis was performed using the Mann–Whitney U test to compare whether there is a difference in the Au accumulation for two treatment groups and p-value < 0.05 was considered significant.

## 3. Results and discussion

### 3.1 Characterization results of SPIO-Au, SPIO-Au-PEG and SPIO-Au-PEG-insulin NPs

The TEM images showed that the synthesized SPIO-Au NPs possess a quasi-spherical shape. The diameters of the SPIO and SPIO-Au NPs were 11.8 nm and 17.5 nm, respectively (Fig. 5). To verify the functionalization process, the hydrodynamic diameter and Zeta-potential of SPIO-Au, SPIO-Au-PEG1, SPIO-Au-PEG2 and SPIO-Au-PEG-insulin NPs were measured from triplicated samples as shown in table 2. The results showed that after the conjugation of SPIO-Au NPs with PEG, the hydrodynamic diameter of first set of SPIO-Au-PEG1 NPs changed from 23.64 to 37.95 nm, while the second set was changed to 76.64 nm. After the further conjugation of SPIO-Au-PEG2 NPs with insulin, the hydrodynamic diameter changed from 76.64 to 121.87 nm. The increase of the hydrodynamic diameter after each step of conjugation is because of the binding of PEG and insulin to the SPIO-Au NP surface, sequentially. After functionalization with PEG, the zeta-potential was changed from −42.30 to −26.43 mV for the first set and −22.67 mV for the second set. Then after conjugation with insulin, the zeta-potential of SPIO-Au-PEG-insulin was further changed from −22.67 to −17.00 mV. These results are in accordance with literature that the PEG and insulin conjugation of AuNPs led to the less-negative zeta potential of NPs and suggest the successful functionalization of SPIO-Au NPs with PEG and insulin [51].

**Table 2.** The zeta potential and hydrodynamic diameter measurement of the NPs

| Particle type | Hydrodynamic diameter (nm) | Zeta-potential (mV) |
| --- | --- | --- |
| SPIO-Au | $23.64 \pm 0.08$ | $-42.30 \pm 0.75$ |
| SPIO-Au-PEG1 | $37.95 \pm 0.34$ | $-26.43 \pm 0.24$ |
| SPIO-Au-PEG2 | $76.64 \pm 0.14$ | $-22.67 \pm 0.22$ |
| SPIO-Au-PEG-insulin | $121.87 \pm 1.17$ | $-17.00 \pm 0.09$ |

**Fig. 5.**
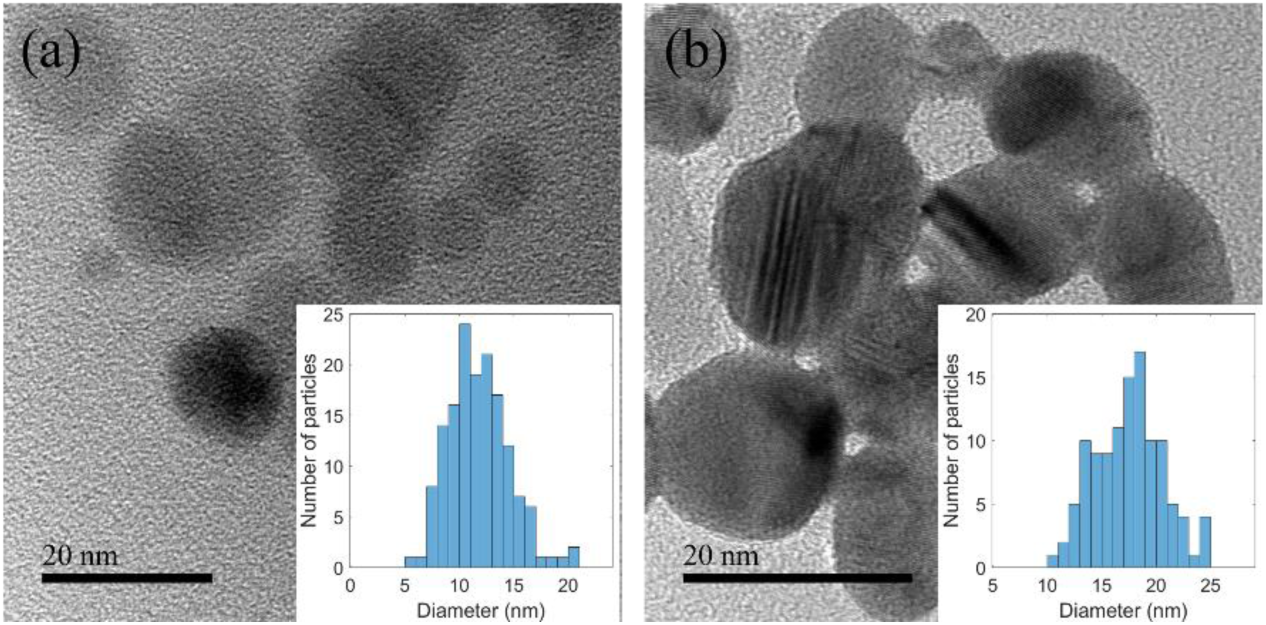
TEM images of (a) SPIO NPs and (b) SPIO-Au NPs. The insets show the size distribution of SPIO and SPIO-Au NPs.

### 3.2 The impact of different magnetic fields on in vivo biodistribution of SPIO-Au-PEG NPs

We first investigated the effect of different magnetic fields (MF-, SMF and DMF) on the accumulation of SPIO-Au-PEG NPs in the brain. The blood and three sections of brain (midbrain, cortex and cerebellum) were collected at 4, 8 and 12 hr after IP injection of SPIO-Au-PEG1 NPs at 30 mg/kg. To eliminate the effect of different absorption rates of SPIO-Au-PEG NPs from the peritoneal cavity to systemic circulation in different mice, the Au level of different brain sections were normalized to the blood. As shown in Fig. 6, the SMF (the average product of the magnetic intensity and its magnetic gradient is 0.74 × 10^13^ A^2^/m^3^) and DMF did not trigger significantly different Au levels in brains for all time points. The average bioavailability from blood to 3 different brain areas are given in table 3 and are calculated as the ratio of the area under the curve (AUC) of the brain area over the blood by plotting mean Au concentrations versus time points. The SMF treatment group shows slightly better bioavailability (4.01%) in the brain compared with MF- (3.68%) and DMF (3.72%). The difference between the SMF and DMF may be due to the periodic changes of the magnetic field intensity and the gradient of the DMF, causing the magnetic force applied on SPIO-Au-PEG NPs to be zero at some time points, unlike SMF which applies the constant magnetic force continually.

**Table 3.**
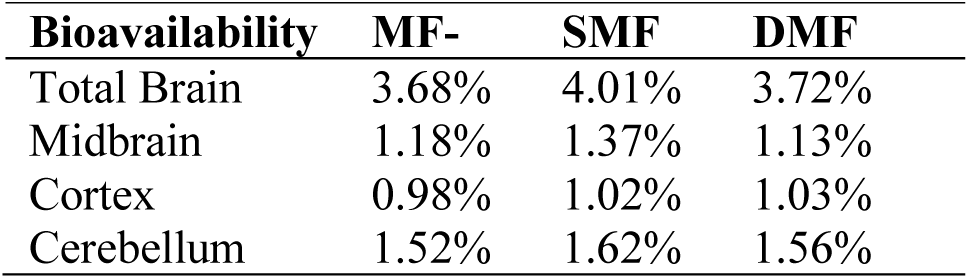
The bioavailability of SPIO-Au-PEG NPs under different magnetic treatments.

| Bioavailability | MF- | SMF | DMF |
| --- | --- | --- | --- |
| Total Brain | 3.68% | 4.01% | 3.72% |
| Midbrain | 1.18% | 1.37% | 1.13% |
| Cortex | 0.98% | 1.02% | 1.03% |
| Cerebellum | 1.52% | 1.62% | 1.56% |

**Fig. 6.**
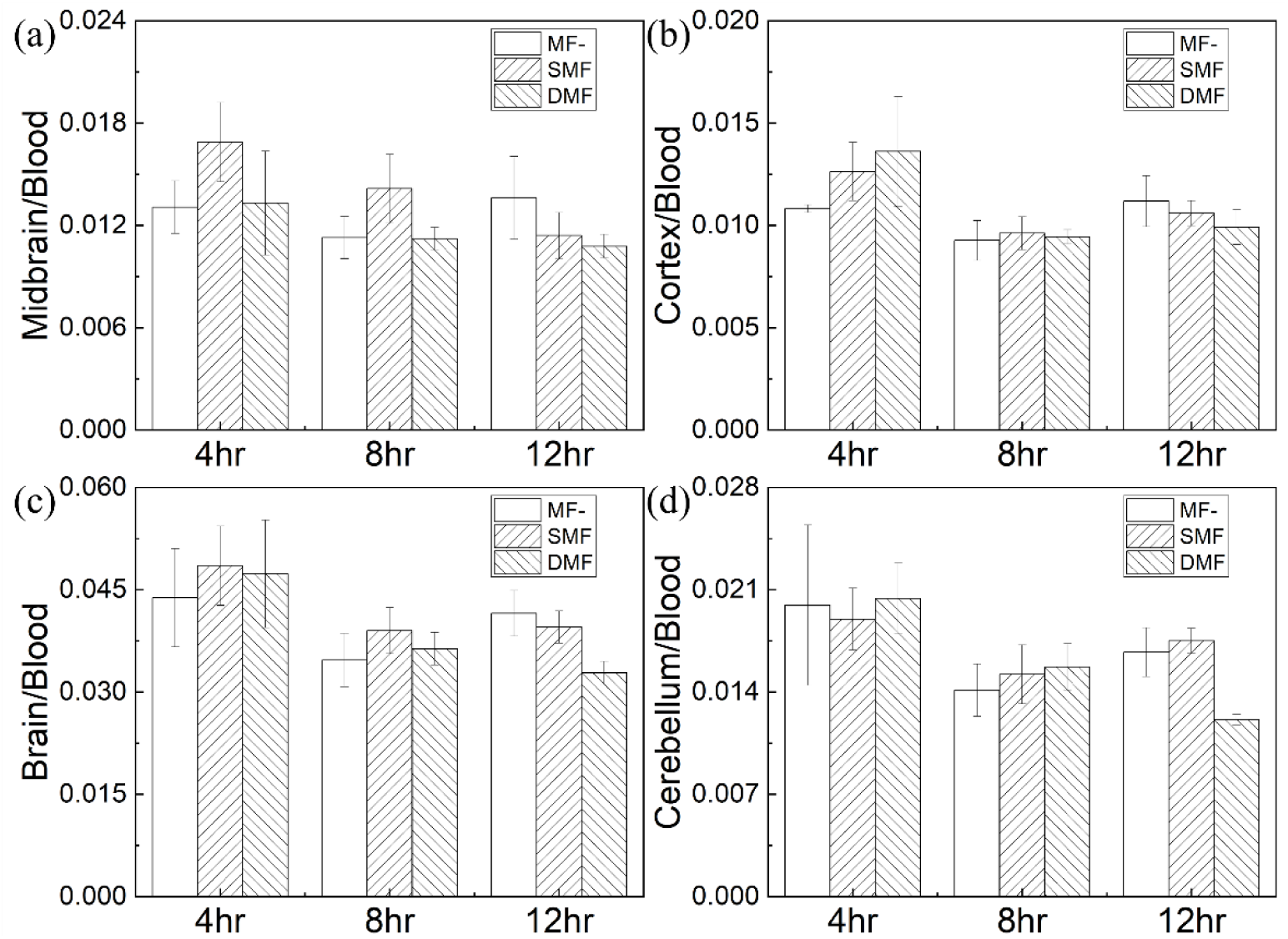
Normalization of Au level in (a) midbrain (b) cortex (c) cerebellum and (d) brain at 4, 8 and 12 hr after a 30 mg/kg IP injection with SPIO-Au-PEG1 NPs under different magnetic treatments (MF-: no magnetic field; SMF: the magnetic field distribution as shown in Fig. 4a; DMF: same with SMF but at a rotation speed of 60 rpm). Error bars represent ± (SEM).

### 3.3 Simulation of brain permeability using advection convection equations

To solve the ODEs that make up the PBPK model, we needed to obtain the brain permeability first. The dynamic behavior of SPIO-Au-PEG NPs in the brain was simulated using the Transport of Diluted Species Module in COMSOL Multiphysics, which solved the advection-convection equations describing the dynamics of SPIO-Au-PEG NPs in the brain. The parameters listed in Table 4 were used in the simulation.

**Table 4.** Parameters used for simulating the behavior of SPIO-Au-PEG NPs in the brain.

| Parameter | Symbol | Value |
| --- | --- | --- |
| SPIO radius<sup>a</sup> | $a$ | 5.4 nm |
| SPIO-Au radius<sup>a</sup> | $R$ | 8.8 nm |
| SPIO-Au-PEG hydrodynamic radius<sup>a</sup> | $R_H$ | 38.3 nm |
| SPIO density<sup>a</sup> | $\rho_{SPIO}$ | 5240 kg/m<sup>3</sup> |
| Au density<sup>a</sup> | $\rho_{Au}$ | 19320 kg/m<sup>3</sup> |
| Initial SPIO-Au NPs concentration<sup>a</sup> | $C_{in}$ | 1 mol/m<sup>3</sup> |
| Magnetic intensity* magnetic gradient<sup>b</sup> | $\nabla(\vert \vec{H}\vert ^2)$ | $1.46 \times 10^{13}$ A<sup>2</sup>/m<sup>3</sup> |
| ICA inflow rate<sup>c</sup> | $Q_{ICA}$ | 0.012 L/h |
| VA inflow rate<sup>c</sup> | $Q_{VA}$ | 0.004 L/h |
| Blood viscosity<sup>c</sup> | $\eta$ | 0.003 Pa.s |
| Temperature<sup>c</sup> | $T$ | 310 K (body temperature) |
| Brownian diffusion coefficient<sup>d</sup> | $D_B$ | $7.27 \times 10^{-12} \text{ m}^2/\text{s}$ |
| Blood cells diffusion coefficient<sup>d</sup> | $D_S$ | $3.92 \times 10^{-10} \text{ m}^2/\text{s}$ |
| Total diffusion coefficient in blood<sup>d</sup> | $D_{total}$ | $3.99 \times 10^{-10} \text{ m}^2/\text{s}$ |
| Diffusion coefficient (in membrane)<sup>d</sup> | $D_{endo}$ | $1.15 \times 10^{-12} \text{ m}^2/\text{s}$ |
| Diffusion coefficient (in tissue)<sup>d</sup> | $D_{tissue}$ | $2.23 \times 10^{-10} \text{ m}^2/\text{s}$ |

<sup>a</sup>values obtained from the NPs characterization.
<sup>b</sup>value obtained from the simulation of the magnetic applicator in COMSOL.
<sup>c</sup>values were from the literature [46].
<sup>d</sup>values were from the equations (6-10).

By solving the advection-convention equations, the concentration of SPIO-Au-PEG NPs in the cerebral blood and brain tissue without magnetic field (MF-) and with the SMF (linear Halbach array of magnets as shown in Fig. 4b) was obtained. Fig. 7a shows the size of the vasculature and the distribution of SPIO-Au-PEG NPs in the CoW at the beginning and after 60 s for MF-(Fig. 7b) and SMF conditions. (Fig. 7c). The concentration-time profile was plotted in Fig. 7d. As shown in Fig. 7d (left), most of SPIO-Au-PEG were quickly washed out by the blood within 1 second. The SMF treatment did not affect the blood concentration of SPIO-Au-PEG within this short time period. As shown in Fig. 7d (right), SMF enhanced the NPs accumulation in the brain tissue: about 8.47% of SPIO-Au-PEG NPs can cross the BBB and stay in the brain tissue for the SMF group, while the value is only 3.36% for MF-treatment group. According to these concentration profiles, the BBB permeability coefficient in the PBPK model was estimated to be 1.32×10^−4^ for MF-condition and 3.51×10^−4^ for SMF condition.

**Fig. 7.**
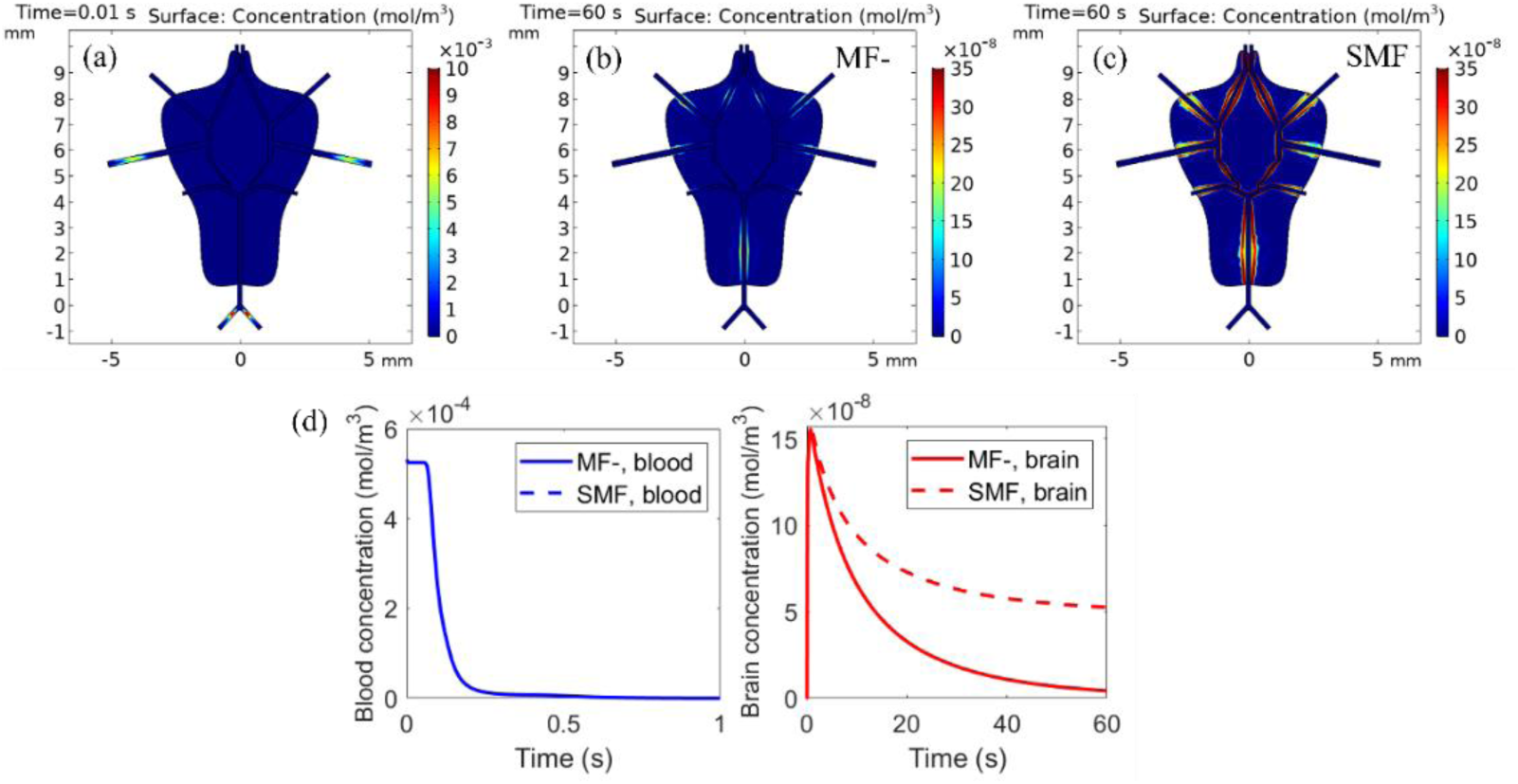
The concentration of SPIO-Au-PEG NPs at t=0.01 (a) and t =60s under the treatment with MF- (b) and with SMF (c); (d) the average concentration in the blood (left) and brain tissue (right). The blue solid line and blue dashed line overlap with each other, indicating that the SMF treatment did not affect the blood concentration of SPIO-Au-PEG NPs.

### 3.4 PBPK model prediction

The brain permeability was incorporated into the PBPK model, while other pharmacokinetic parameters of SPIO-Au-PEG NPs were first estimated based on the similar PBPK model [40], which predicted the concentration of PEG coated AuNPs (13-20 nm, diameter) in the mice (0.02 kg, weight). These parameters were obtained by fitting the unknown parameters against the experimental data using the Nelder-Mead method. We further optimized the parameters by manually adjusting the parameters iteratively with the measured *in vivo* data for the MF-group until a reasonable match between the *in vivo* data and the PBPK model prediction was reached. The final parameters were listed in table 5. The manual fitting approach has multiple advantages over the computational-based approach in the development of the PBPK model for SPIO-Au-PEG NPs in this study. Firstly, there were many unknown parameters that had to be estimated based on a single dataset, so the estimated values undoubtedly had a large deviation. Secondly, due to the sparseness of the *in vivo* data (limited time point data) which led to a great uncertainty in the concentration-time profiles, it was not feasible to directly use a computational-based approach for parameter estimation. Thirdly, by using the manual fitting approach, some observations from the *in vivo* study could be considered and incorporated into the model calibration, which made the parameter estimation more reasonable.

**Table 5.** Pharmacokinetic related parameters of the PBPK model

| Parameter | Liver | Spleen | Kidneys | Lungs | Brain | Remainder |
| --- | --- | --- | --- | --- | --- | --- |
| $P_o$ (unitless) | 0.08 | 0.15 | 0.15 | 0.15 | 0.15 | 0.15 |
| $PAC_o$ (unitless) | $10^{-3}$ | $10^{-3}$ | $10^{-3}$ | $10^{-3}$ | $1.32 \times 10^{-4}$ (MF-)<br>$3.51 \times 10^{-4}$ (SMF) | $10^{-6}$ |
| $K_{max_o}$ (/h) | 4 | 10 | 0.1 | 0.1 | NA | NA |
| $K_{half_o}$ (h) | 24 | 24 | 24 | 24 | NA | NA |
| $n_o$ (unitless) | 0.1 | 0.1 | 0.1 | 2 | NA | NA |
| $K_{release_o}$ (/h) | 0.0075 | 0.003 | 0.01 | 0.005 | NA | NA |
| $K_{bile}$ or $K_{urine}$ (L/h) | 0.0012 | NA | 0.00012 | NA | NA | NA |

As shown in Fig. 8a, the simulated data matches well with the measured *in vivo* data, indicating that the model was well calibrated to accurately predict the concentrations in the blood, brain, liver and lung at all time points for the MF-groups. Unlike IV injection in which NPs directly enter into systemic circulation and rapidly reach a peak, the IP injected NPs undergo an absorption process before entering systemic circulation. Specifically, after IP injection, the amount of Au in the blood slowly increases as NPs were constantly absorbed from the peritoneal cavity. Following the distribution of SPIO-Au-PEG NPs to the organs, the concentration of Au in the blood decreases with the peak level occurring around 6 hr post injection. The amount of Au in the brain has a similar trend to that in the blood. However, the amount in the liver and lung increases quickly from 0 to 16 hr, indicating the saturation point is still not reached. A similar trend was also observed in PEG coated AuNPs from 0 to 30 hr [24]. The overall regression coefficient (goodness of fit) between measured and predicted data is R^2^ = 0.936 as shown in Fig. 8b.

**Fig. 8.**
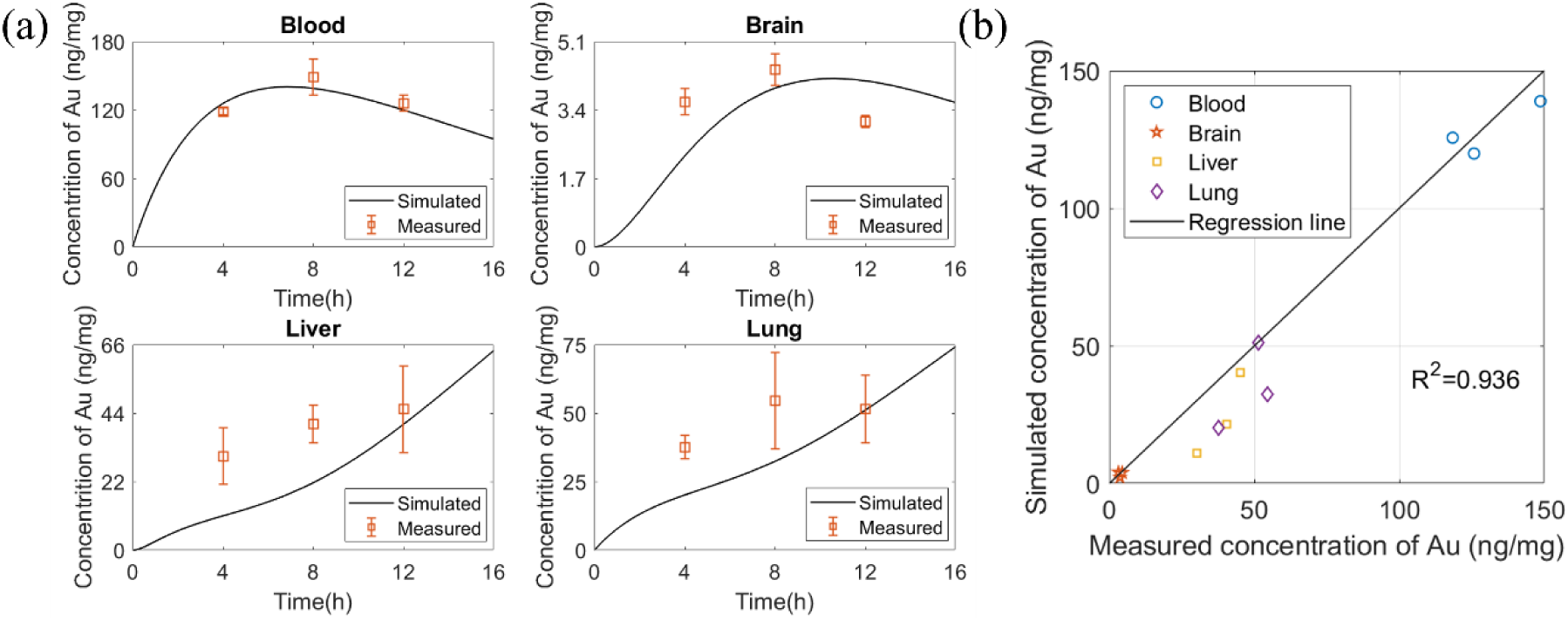
(a) Amount of Au per organ weight with MF-predicted by the PBPK model versus measured data in blood, liver, lung, and brain. Error bars show the standard deviation of measured data; (b) Goodness-of-fit plot of the linear regression analysis of model predictions and measured data (R^2^ = 0.936).

Next, we employed the PBPK model to study the pharmacokinetics of SPIO-Au-PEG NPs under the exposure of SMF. As shown in Fig. 9a, by comparing the simulated and measured data, the PBPK model properly predicted the dynamics of NPs in the blood and brain, with a slight (2-fold) underestimation of the concentrations in the liver and lung. It is possible that after IP injection, partial SPIO-Au-PEG NPs are transported directly to the liver via the portal vein before they enter systemic circulation [52], causing the higher measured concentration of Au in liver at all time points and accounting for the underestimation in the model. The overall regression coefficient between measured and predicted data is R^2^ = 0.825 as shown in Fig. 9b.

**Fig. 9.**
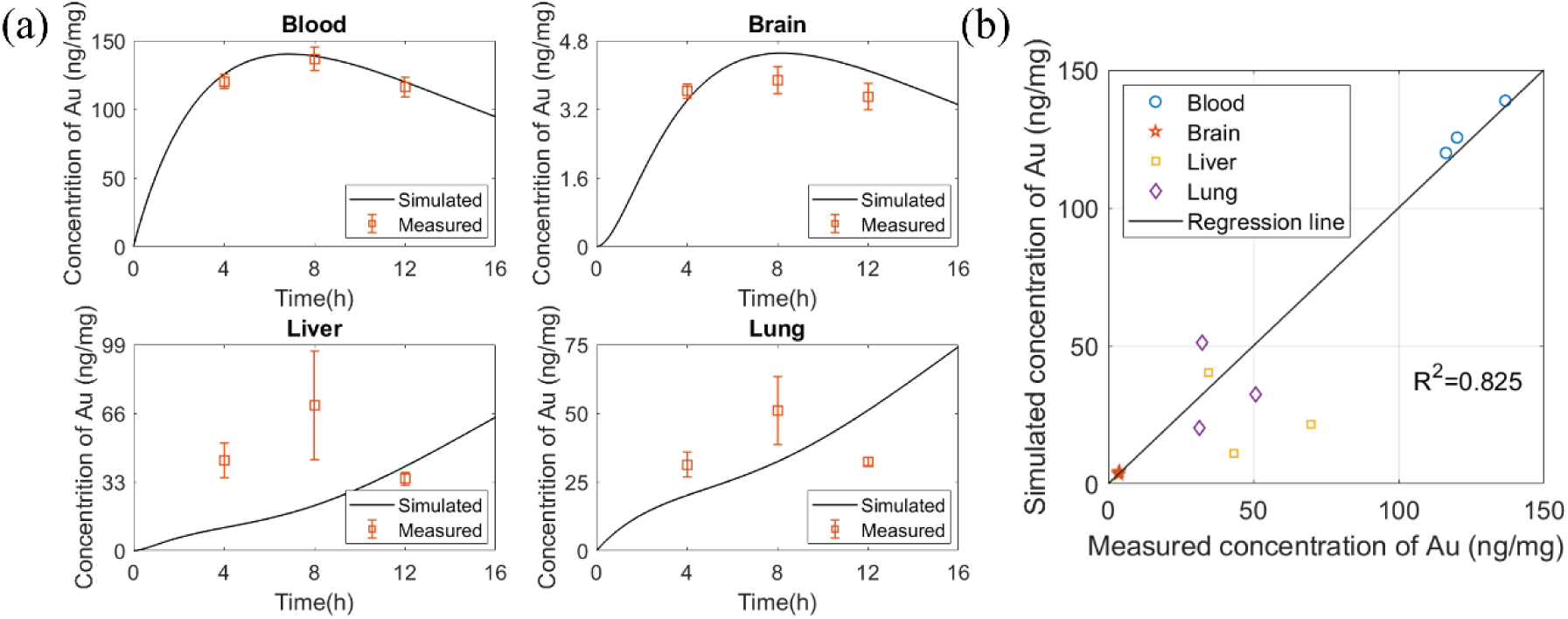
(a) Amount of Au per organ weight with SMF predicted by the PBPK model versus measured data in blood, liver, lung, and brain. Error bars show the SEM of measured data; (b) Goodness-of-fit plot of the linear regression analysis of model predictions and measured data (R^2^ = 0.825).

### 3.5 The impact of insulin on biodistribution of SPIO-Au-PEG NPs

To further enhance the BBB crossing, we investigated the effect of insulin modification of the SPIO-Au-PEG NPs on the *in vivo* biodistribution. Mice were treated with either (1) insulin- or (2) insulin+ without exposure to magnetic field. The blood, midbrain, cortex and cerebellum were collected at 4, 8 and 12 hr after NPs injection. Fig. 10a-d shows the Au level in different brain areas normalized to the blood. It is observed that at the time point of 4 hr, the insulin added group shows a significantly higher Au accumulation in the midbrain (2.1 times) and cortex (1.6 times) compared to the control group (insulin-). While in the cerebellum, this enhanced Au accumulation by insulin conjugation is observed at 12 hr. The results of total brain reflect that the insulin added groups show higher brain Au accumulation at 12 hr post injection. With the conjugation of SPIO-Au-PEG with insulin, the bioavailability of SPIO-Au-PEG NPs from blood to brain was increased from 2.86% to 3.56% (an improvement by 24.47%). The results indicated the accumulation of SPIO-Au NPs in different brain sections were time-related and the addition of insulin enhanced the bioaccumulation of SPIO-Au NPs in the brain. It is noticeable that the control group has a brain bioavailability of 2.86% which is smaller than the control group (3.68%) in section 3.2. The reason is likely that to enable the insulin conjugation, about 15% of the SH-mPEG (5 kDa) is replaced by SH-PEG-COOH, which has smaller molecular weight (3.4 kDa), during the insulin conjugation process. This may cause the slightly lower blood circulation time [37] which may induce the lower brain bioavailability because of the slightly lower chance of NPs interacting with the BBB.

**Fig. 10.**
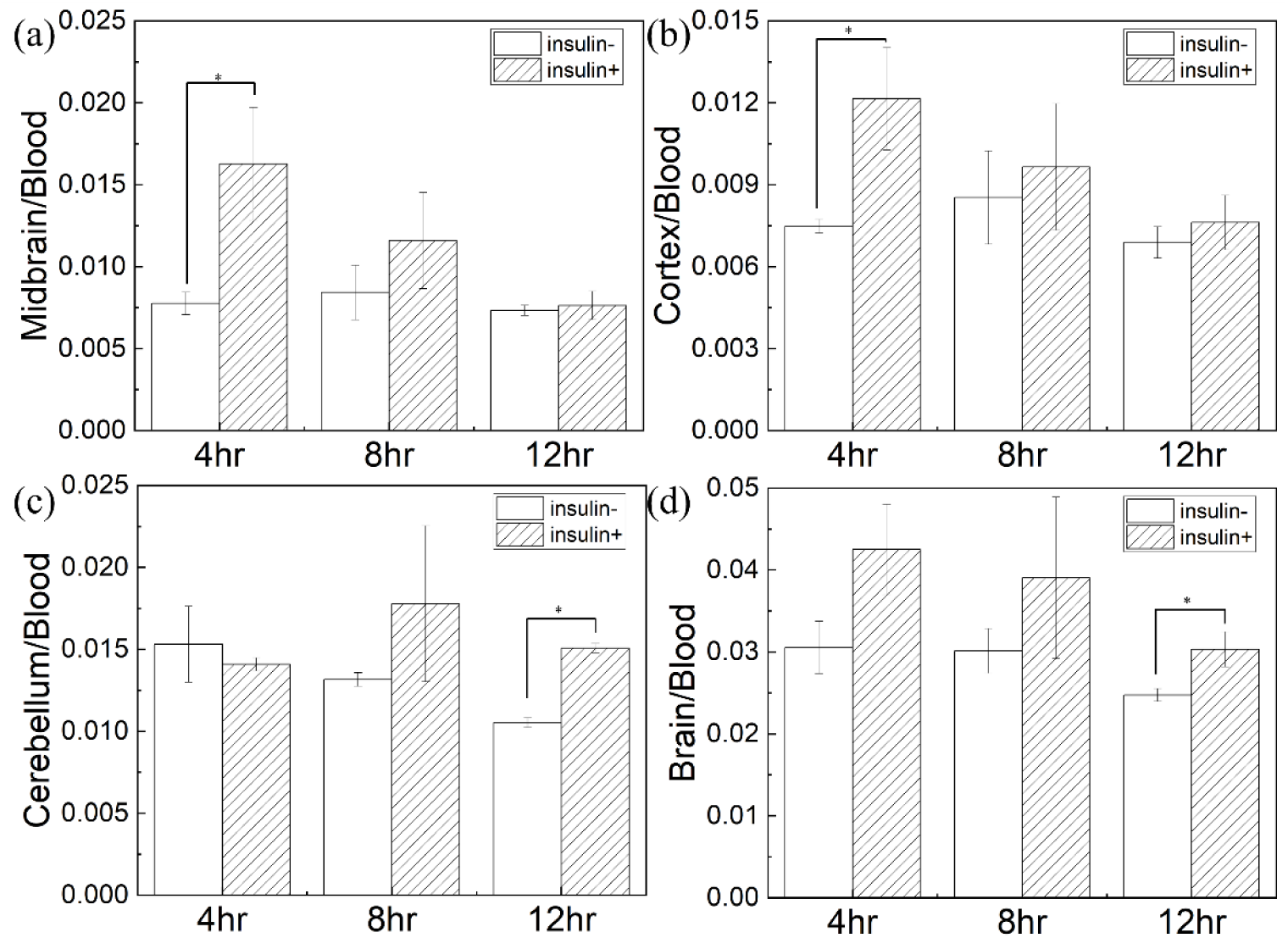
Normalization of Au level in (a) midbrain (b) cortex (c) cerebellum and (d) total brain to the blood at 4, 8, and 12 hr after IP injection with NPs at the dosage of 30 mg/kg without magnetic treatment (insulin-: injected with SPIO-Au-PEG2 NPs; insulin+: injected with SPIO-Au-PEG-insulin NPs). Error bars represent ± (SEM), p-values were calculated from nonnormal distributions using the Mann–Whitney U test (*: p < 0.05).

The static magnetic applicator was then modified to increase the magnetic field intensity and its gradient (∼1.75 times) as shown in Fig. 4b. To study the effect of the enhanced SMF field on the accumulation of SPIO-Au-PEG-insulin NPs, the blood, midbrain, cortex and cerebellum were collected at 2, 4, 8 and 12 hr after IP injection of SPIO-Au-PEG-insulin NPs at 30 mg/kg with MF- and SMF. The normalized Au level of different brain areas to the blood are demonstrated in Fig. 11a to d. The results show that at the time point of 2 hr, the SMF significantly enhances the Au level in midbrain, cortex, cerebellum and total brain compared with the non-MF condition. Additionally, it is observed that at this time point the SMF treatment increases the Au accumulation up to 1.46 times for the cortex and 1.54 times for the cerebellum, which is higher than the Au accumulation for the midbrain (1.33 times) in accordance with the magnetic field distribution in which the cortex and cerebellum were exposed to a stronger magnetic field compared to the midbrain area. The bioavailability of SPIO-Au-PEG-insulin NPs from blood to brain for MF- and SMF groups, are 3.58% and 3.72% respectively (an improvement by 3.91%). The results indicate that the treatment with the enhanced SMF increases the accumulation of SPIO-Au-PEG-insulin NPs in the brain compared to the control group at the time point of 2 hr, suggesting the brain targeting effect of SMF. However, for the longer time periods, this effect is not observed. Also, as shown in supplementary material Fig. S1a-b, the Au amount in the liver and lung did not show much difference between treatment groups, suggesting that the SMF treatment does not affect the biodistribution of SPIO-Au-PEG NPs in other tissues except for the brain.

**Fig. 11.**
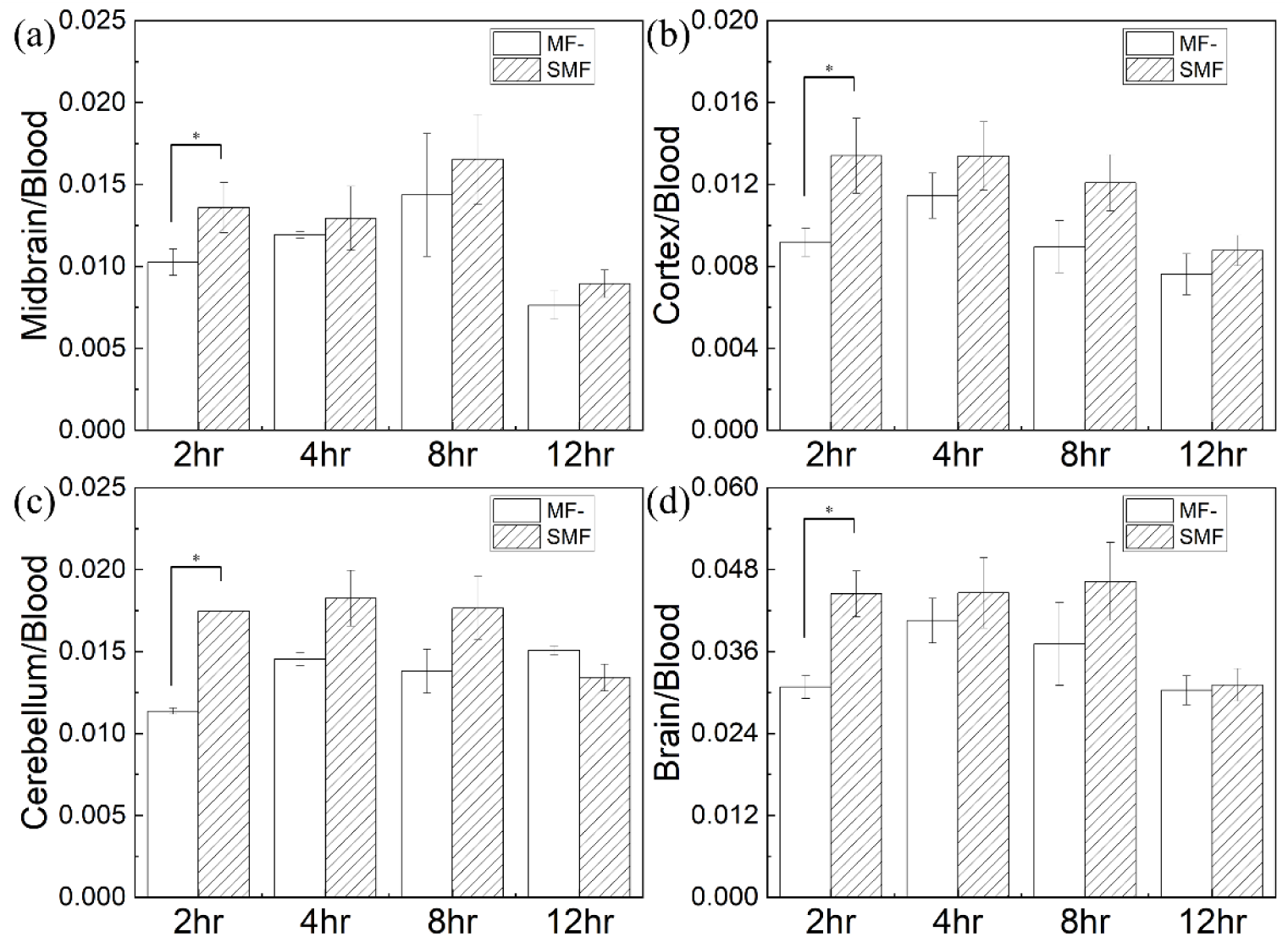
Normalization of Au level in (a) midbrain (b) cortex (c) cerebellum and (d) total brain to the blood at 2, 4, 8 and 12 hr after IP injection of SPIO-Au-PEG-insulin NPs at the dose of 30 mg/kg (MF-: no magnetic applicator; SMF: the magnetic field distribution as shown in Fig. 4b). Error bars represent ± (SEM), p-values were calculated from nonnormal distributions using the Mann–Whitney U test (*: p < 0.05).

In this study, we only employed the PBPK model for SPIO-Au-PEG NPs. The reasons are as follows. (1) The objective of the PBPK model is to provide a quantitative tool to simulate the behavior of MNPs in the brain under an external SMF. By using the PBPK model to predict the biodistribution of SPIO-Au-PEG NPs under the exposure of SMF, we demonstrated that this model which considered the whole-body circulation and the complexity of the BBB is a feasible approach. (2) The pharmacokinetic parameters in the PBPK model are NP-dependent values. Conjugation of SPIO-Au-PEG with insulin changes the parameters, such as the endocytosis rate, making the PBPK model more complicated to develop with the limited dataset.

## 4. Conclusions

In this study, we have carried out a comprehensive analysis from theoretical, numerical and *in vivo* perspectives to understand and quantify the dynamic BBB crossing behavior of SPIO-Au-PEG NPs. We successfully developed a theoretical PBPK model for predicting the *in vivo* biodistribution of SPIO-Au-PEG NPs in mice after IP injection. Specifically, the brain permeabilities of SPIO-Au-PEG NPs under different conditions (no magnetic field and SMF) were calculated by obtaining their concentration from the advection-diffusion equations. This PBPK model has been calibrated and validated using our *in vivo* results, indicating its robust predictive capability. From *in vivo* results we also demonstrated the conjugation of SPIO-Au-PEG NPs with insulin facilitates the BBB crossing and increases the bioavailability in the brain from 2.86% to 3.56%. Furthermore, the bioavailability can be improved to 3.72% under exposure to SMF.

The PBPK model provides insights into the pharmacokinetics of SPIO-Au-PEG NPs in *vivo*; thus, it can reduce the use of living animals for testing and significantly decreases the experimental cost at this stage for studying the magnetic targeting efficiency of MNPs. This theoretical modeling, numerical simulation and *in vivo* validation lays a solid foundation towards non-invasive brain therapeutics with maximal accuracy and minimal side effects.

The simulation and *in vivo* measurements show that magnetic stimulation can enhance BBB crossing; however, the small size of the magnetic core limits the magnetic targeting efficiency. As the radius of the magnetic core increases, the magnetic force applied on it will increase by the cube of the increasing factor. To overcome the low driving magnetic force applied on SPIO-Au NPs and increase the BBB crossing efficiency, in future work, we will develop a biodegradable hydrogel micro-swimmer containing SPIO-Au NPs with a stronger magnetic response. Additionally, this PBPK model will be further modified and employed to study its targeting efficiency.

## Acknowledgements

This work was kindly supported by the United States National Science Foundation (NSF) (Award # CMMI 1851635, Y.W.; Award # ECCS 2021081, Y.W. and S.E.).

